# Gene-based association analysis identified 190 genes with polymorphisms affecting neuroticism

**DOI:** 10.1101/2020.08.06.240721

**Authors:** Nadezhda M. Belonogova, Irina V. Zorkoltseva, Yakov A. Tsepilov, Tatiana I. Axenovich

## Abstract

Recent genome-wide studies have reported about 600 genes potentially influencing neuroticism. Little is known about the mechanisms of their action. Here, we aimed to conduct a more detailed analysis of genes whose polymorphisms can regulate the level of neuroticism. Using UK Biobank-based GWAS summary statistics, we performed a gene-based association analysis using four sets of genetic variants within a gene differing in their protein coding properties. To guard against the influence of strong GWAS signals outside the gene, we used the specially designed procedure. As a result, we identified 190 genes associated with neuroticism due to their polymorphisms. Thirty eight of these genes were novel. Within all genes identified, we distinguished two slightly overlapping groups comprising genes that demonstrated association when using protein-coding and non-coding SNPs. Many genes from the first group included potentially pathogenic variants. For some genes from the second group, we found evidence of pleiotropy with gene expression. We demonstrated that the association of almost two hundred known genes could be inflated by the GWAS signals outside the gene. Using bioinformatics analysis, we prioritized the neuroticism genes and showed that the genes influencing the trait by their polymorphisms are the most appropriate candidate genes.

## Introduction

Neuroticism is a relatively stable ^1^ and heritable personality trait ^2^, which is an important risk factor for psychiatric disorders ^3,4^. The strong genetic correlation between neuroticism and mental health ^5–7^ suggests that neuroticism may represent an intermediate phenotype, which is influenced by risk genes more directly than psychiatric diseases ^8^. This implies that exploring the genetic contribution to differences in neuroticism can help understand the genetic architecture of psychiatric disorders.

Until recently, just a few GWAS loci for neuroticism were identified ^9–11^. Several possible candidate genes have been suggested at these loci, including *GRIK3*, *KLHL2*, *CRHR1*, *MAPT*, *CELF4*, *CADM2*, *LINGO2* and *EP300* ^12^. Some of these genes have been reported to be associated with depression, autism, Parkinson’s disease, schizophrenia and other psychiatric disorders.

Great progress in the genetic dissection of neuroticism was made in 2018, when 170 ^13^ and 116 ^6^ independent SNPs associated with neuroticism were identified using large samples of 449 484 and 329 821 people, respectively. These SNPs marked 157 loci, 64 of which were identified in both studies. The majority of the lead SNPs were located in intronic and intergenic regions, and only a few SNPs were coding. With the help of positional mapping, Nagel et al. ^13^ identified 283 genes potentially influencing neuroticism and located at independently associated loci. The list of genes for neuroticism was expanded using genome-wide gene-based association and expression quantitative trait locus (eQTL) analyses, and chromatin interaction mapping. In total, 599 genes were included in this list, 50 of them were identified by all four methods. It was demonstrated that these genes were predominantly expressed in six brain tissue types and were associated with seven Gene Ontology (GO) gene sets including neurogenesis, neuron differentiation, behavioral response to cocaine, and axon part ^13^.

In our study, we focus on further identification of genes that are associated with neuroticism and their detailed functional analysis. We concentrated on the genes whose polymorphisms are directly responsible for differences in the individual phenotype. Polymorphisms within the genes can modify the structure of the corresponding proteins due to amino acid substitutions or alternative splicing. Polymorphisms in the non-coding intragenic regions can regulate transcription and translation of these genes, protein complex formation or posttranslational modifications ^14,15^. Such modifications may tilt the physiological balance from healthy to diseased state, resulting, for example, in bipolar affective disorder or Alzheimer’s disease ^16^. Altogether, within-gene association signals are usually easy to interpret and less probable to be false-positives.

Using summary statistics obtained from UK Biobank data, we performed a gene-based association analysis investigating several sets of genetic variants that differ by their protein-coding properties. We used a procedure that we called ‘polygene pruning’ to guard against the influence of strong GWAS signals outside the gene. As a result, we have identified 190 genes associated with neuroticism, both known and novel. We performed a bioinformatics analysis to compare the biological functions of the known genes confirmed and non-confirmed in our study, and the biological functions of the known and novel genes, and to check if the identified genes can be considered as true gene candidates for neuroticism.

## Results

### The gene-based analysis of neuroticism

The strategy of our study and the main results are shown in Figure 1. The Manhattan plot for the association signals obtained before and after polygene pruning is presented in Figure 2a. The genes identified under the different scenarios after polygene pruning are shown in Supplementary Tables 1-4. The full list of 190 genes identified under at least one scenario is presented in Supplementary Table 5.

**Fig. 1.**
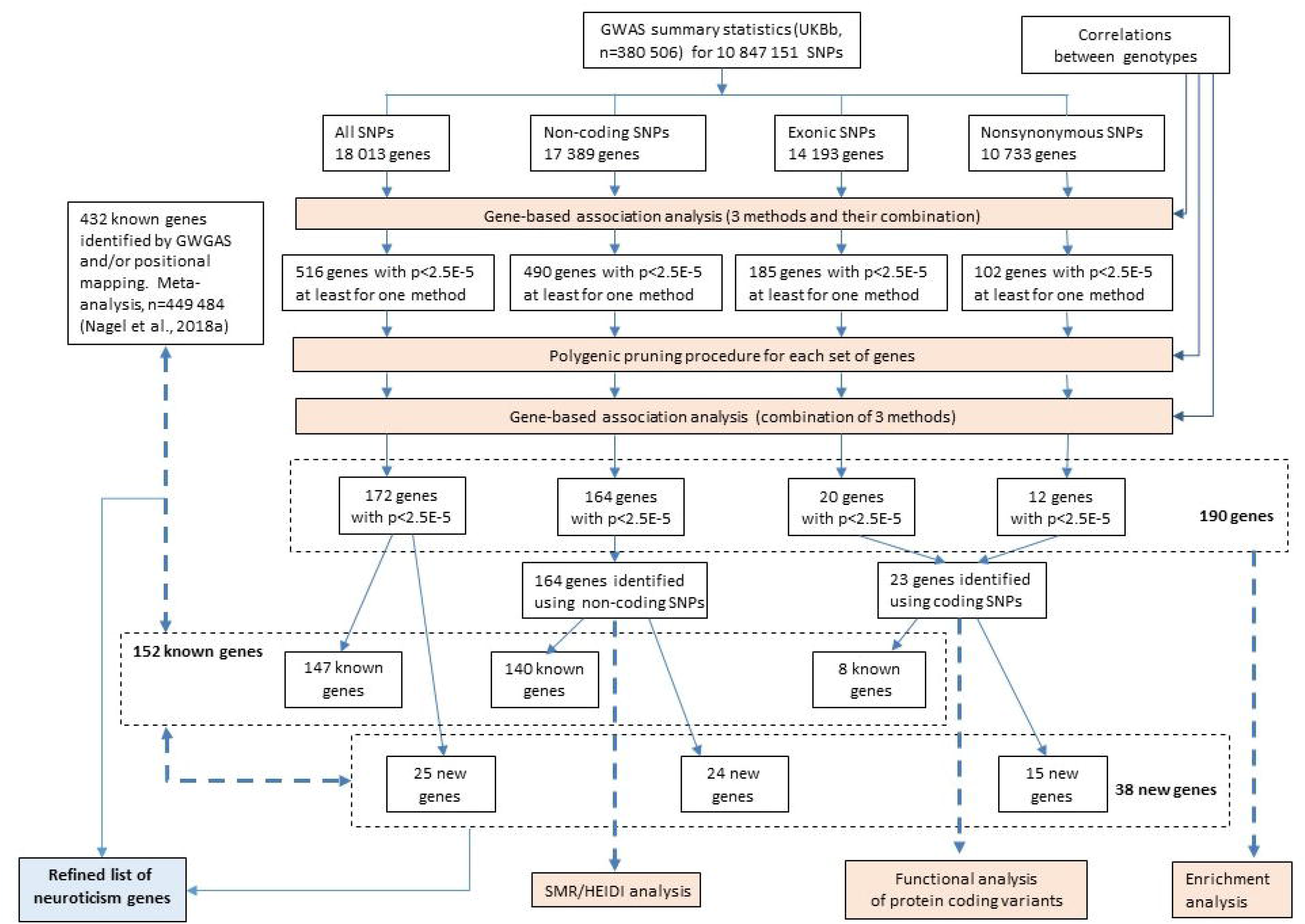
Workflow schematic. GWAS summary statistics and correlations between genotypes were used as input data. Each set of SNPs (all, non-coding, exonic, nonsynonymous) was analyzed separately. The first step of our study is the gene-based association analyses performed using the SKAT-O, PCA, ACAT methods and their results in combination. Next, to guard against the influence of strong GWAS signals outside the gene, we performed polygene pruning for genes with p-value < 2.5×10^−5^ demonstrated at least for one of the gene-based methods. After pruning, we repeated the gene-based analysis using the remaining SNPs. The genes identified using each set of SNPs were subdivided into known and novel. All the known genes confirmed in our study were compared with the rest of the known genes for their functional properties. We also compared the new and known genes identified in our study. For all identified genes, we performed an enrichment analysis; for the genes identified using protein-coding SNPs, a functional analysis of protein-coding variants was performed; for the genes identified using non-coding SNPs, SMR/HEIDI analysis was performed. Bold blue arrows show the gene groups being compared and gene groups being under *in-silico* analyses.

**Figure 2.**
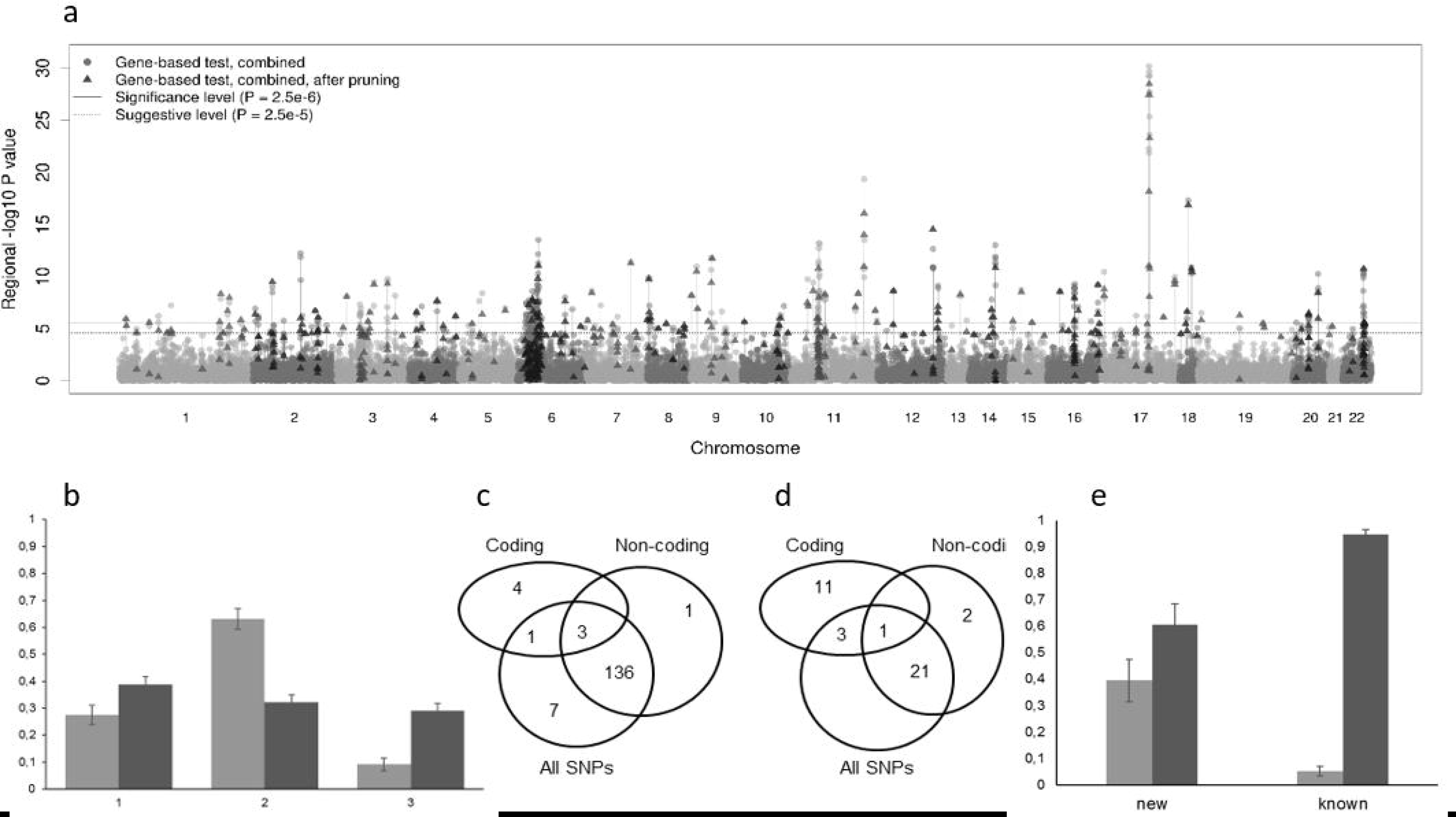
The results of gene-based association analysis. **a.** Manhattan plot showing the −log__10__ transformed p-value of each gene. **b**. Proportion of genes identified by gene-based association analysis only (1), both gene-based association analysis and position mapping (2) and positional mapping only (3) among confirmed (light gray) and non-confirmed (dark gray) genes. **c**. Venn diagram showing the overlap of known genes identified using different sets of SNPs. **d.** Venn diagram showing the overlap of new genes identified using different sets of SNPs. **e**. Proportion of genes demonstrating associations using non-coding (dark gray) and protein-coding SNP sets (light gray) in the new and known genes.

### Comparison with known genes

From among 599 previously suggested neuroticism genes from Supplementary Table 15 in a work by Nagel et al. ^13^, we took a subset including protein-coding genes whose effect on neuroticism can be explained by the SNPs located within these genes. Such genes had been previously identified by gene-based association analysis or/and positional mapping. We added two genes, *PLCL2* and *ARHGAP15*, identified by eQTL analysis because their expression was controlled by SNPs located within these genes. We defined this subset of 432 genes as a list of known genes (Supplementary Table 6).

From among 432 known genes, 152 overlapped with genes identified in our study and 280 did not. From among 280 non-overlapping genes, 190 demonstrated the gene-based association with neuroticism before polygene pruning and lost association after them. From among the overlapping genes, two thirds were included in the list of the known genes by both gene-based analysis and positional mapping, while this proportion was one-third for the non-overlapping genes (Figure 2b). The reverse was observed for the proportion of genes included in the list by positional mapping (Figure 2b).

Then we compared the known and new genes identified in our study. Among 152 known genes, eight were identified using protein coding SNPs; 140, using non-coding variants (three were identified using both coding and non-coding SNP sets); and seven genes using only the all-variants scenario (Figures 1 and 2c, e). From among 38 new genes, 15 were identified using protein-coding SNP sets and 24, using non-coding variants (one gene was identified using both coding and non-coding SNP sets) (Figures 1 and 2d, e).

We have identified 23 genes using protein-coding SNP sets. Three of them demonstrated association when only nonsynonymous SNPs were used. Nine were associated when using both nonsynonymous and exonic variation, with seven of them having the lowest p-values when using nonsynonymous variation only (Supplementary Tables 3 and 4).

### Enrichment analysis of 190 identified genes

Analysis of differentially expressed genes showed a strong enrichment of genes expressed in the different brain structures (Figure 3a and Supplementary Table 7). The most significant enrichment was demonstrated for the following brain tissues: the hypothalamus, cerebellar hemisphere, cerebellum, frontal cortex (BA9), cortex, anterior cingulate cortex (BA24), nucleus accumbens basal ganglia.

**Figure 3.**
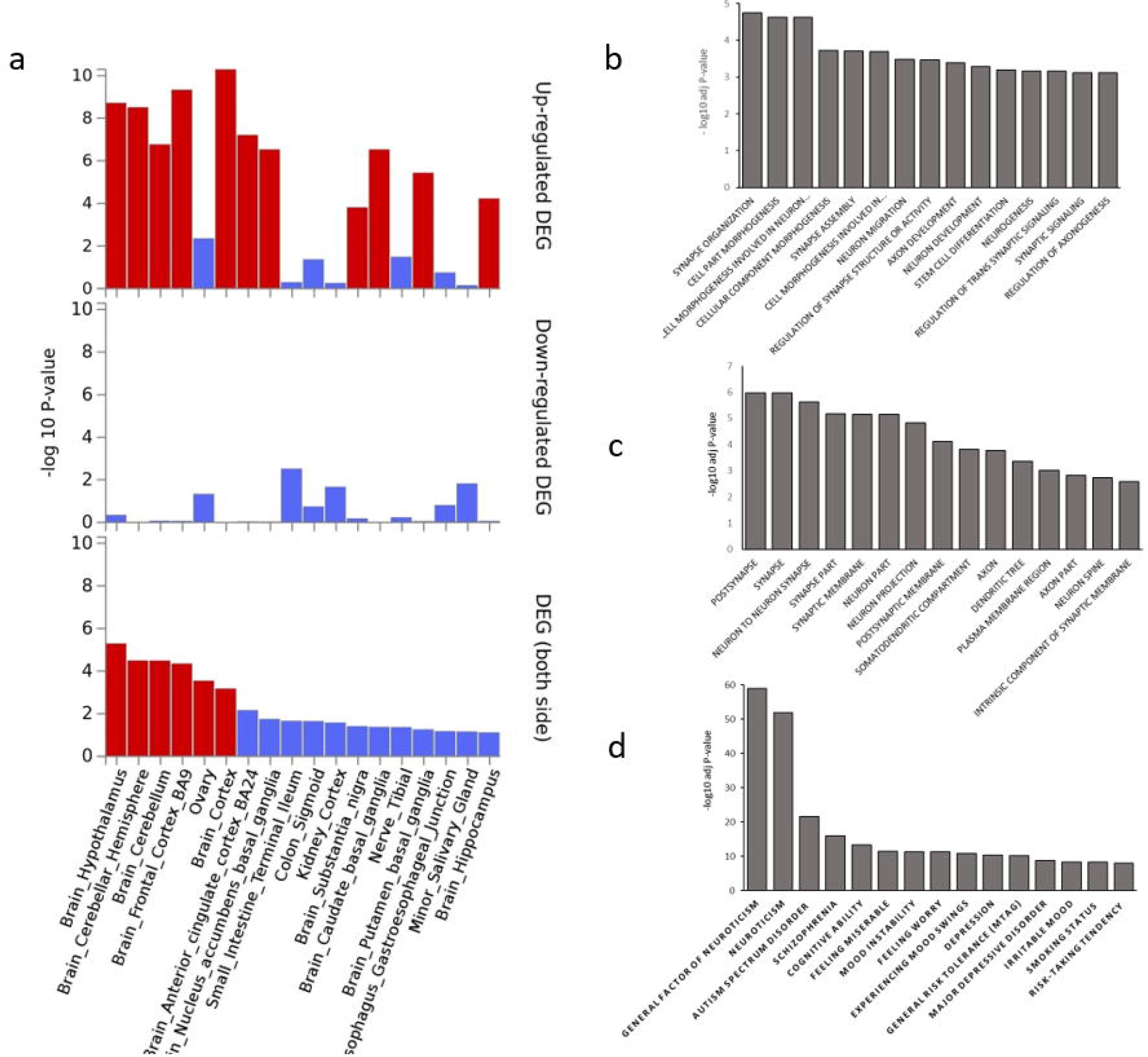
Enrichment of 190 genes identified by gene-based association analysis in different gene sets. **a.** Tissue enrichment for differential expression. **b.** Enrichment in GO biological processes. **c.** Enrichment in GO cell components. **d.** Representation of genes in the GWAS catalog.

To assess whether the identified genes converge on shared biological functions or pathways, we used FUMA for an enrichment test in the GO biological processes and GO cell components. The gene sets with an adjusted p-value ≤ 0.05 are shown in Supplementary Table 8. We detected 58 significantly enriched GO terms: 36 in GO biological processes and 22 in GO cellular components. The top 15 results are shown in Figures 3b and 3c, respectively. Among the biological processes with the most significant enrichment were synapse organization, cell morphogenesis involved in neuron differentiation, synapse assembly, neuron migration, regulation of synapse structure or activity, axon and neuron development, neurogenesis. For cellular component categories, the most significant findings were those for postsynapse, synapse, neuron-to-neuron synapse, synapse part, neuron part, neuron projection, and the others.

We estimated the overrepresentation of the identified genes in the GWAS catalog. The most represented gene sets were associated with psychiatric, cognitive and behavioral traits such as neuroticism, general factor of neuroticism, autism spectrum disorder, schizophrenia, cognitive ability, mood instability, feeling worry, and depression (Figure 3d).

### Functional analysis of protein coding variants

For 23 genes demonstrating association with neuroticism when using protein-coding SNP sets, we selected 76 SNPs with the p-value < 0.05 (Supplementary Table 9). The full description of the effect predicted for SNPs by VEP and FATHMM-XF is presented in Supplementary Tables 10 and Eighteen SNPs in 16 genes were identified as potentially pathogenic variants in accordance with at least one algorithm (Supplementary Table 12). One variant, rs9267835, was defined as pathogenic by all four algorithms.

### SMR and HEIDI analysis of non-coding variants

SMR analysis followed by the HEIDI test provided strong evidence for the pleiotropy of the polymorphisms within fourteen genes with the expression of 25 unique genes in two tissues: whole and peripheral blood (Supplementary Table 13). For eight genes, we detected significant pleiotropic signals with the expression of more than one gene at the same locus. For polymorphisms within six genes (tagged by intronic variants rs75614054, rs11975, rs10896636, rs10005233, rs240769 and rs3996325), we found evidence of pleiotropy with the expression of the same genes where these SNPs were located (*PTCH1, FAM120A, ZDHHC5, SNCA, ASCC3* and *MAD1L1,* respectively). For the polymorphisms in other eight genes, we found evidence of pleiotropy with the transcription of adjacent genes.

## Discussion

In the present study, we have proposed and applied a new approach for detection of genes associated with neuroticism. We have identified 190 genes associated with neuroticism due to their polymorphisms. Thirty eight of these genes were novel.

Our approach is different from those previously applied to study neuroticism in three distinctive ways. First, we used gene-based association analysis to identify neuroticism genes. Due to the simultaneous consideration of a set of genetic variants within a gene, gene-based association analysis has increased power for identification of common and rare genetic variants ^17,18^. Given that, we have identified 38 new neuroticism genes.

Secondly, we used several different sets of SNPs representing different hypotheses of the polymorphism function mediating the association. This allowed us to distinguish two slightly overlapping groups among the associated genes. The first group is the genes whose association is the consequence of the changes in their regulatory properties. The second group is the genes whose association is the consequence of the changes in the structure of the encoded proteins.

Finally, we used the polygene pruning procedure. The aim of this procedure was to reduce the influence of strong association signals located outside the gene. Initially, we identified 459 genes and only 190 remained in the list of the neuroticism genes after polygene pruning. Therefore, we can conclude that the high proportion of gene-based association signals initially obtained in our study was inflated by strong GWAS signals outside the genes. We suggested that this conclusion can be applied, to some extent, to the 432 known neuroticism genes. If so, the genes that were confirmed in our study can be expected to differ in their functions from the genes that were not confirmed. We compared the enrichment of these two groups of genes in GO biological processes and cellular components and found differences between these groups. For GO biological processes, the top results for the confirmed genes included the processes occurring in the nervous system (Supplementary Table 8), while for the non-confirmed ones, the top results were obtained for less specialized processes, such as protein DNA complex subunit organization, chromatin assembly or disassembly, nucleosome organization, cellular protein containing complex assembly chromosome and chromatin organization (Supplementary Table 14). Similar results were obtained using GO cellular components (Supplementary Tables 8 and 14). Two thirds of the confirmed genes were included in the list of the known genes by both gene-based and positional analyses. For the non-confirmed genes, this proportion was one-third. These results have led us to conclude that the known genes confirmed in our study and associated with neuroticism due to their polymorphisms are more appropriate candidate genes than the non-confirmed genes.

Among the genes we have identified were both known and new. These two groups differed in the proportion of genes identified using protein-coding SNPs: 39% (15/38) and 5% (8/152) for the new and the known genes, respectively. This difference can be explained by a high impact of GWAS in the search of common polymorphism associations. The common variants identified by this method are more frequent at intergenic and intronic loci than in protein-coding ones. Due to it, only a low proportion of the known genes was confirmed in our study using coding SNPs. Four (*CRHR1, NOTCH4, NOS1* and *AGBL1*) of eight such genes included common variants with p-value < 5×10^−8^.

Only three (*C12orf49, RPP21* and *TRIM39-RPP21*) of the 15 new genes included such SNPs. The same situation was observed for the genes identified using non-coding SNPs: 72 of 137 known genes and only one of 23 new genes included significantly associated common variants. This confirms that a significant number of the known genes was identified due to strong association signals of common variants. The proportion of the new genes with such variants is rather low. The majority of the new genes have been described as genes associated with intelligence, education attainment, cognitive performance, well-being spectrum, alcohol dependence, worry, anorexia nervosa, autism spectrum disorder, schizophrenia, depression and Parkinson’s disease (Supplementary Table 15). Taking into account a high genetic correlation of neuroticism with mental health ^6,13,19,20^, the new genes identified in our study appear to be reasonable candidates for neuroticism genes. Therefore, both known and novel genes identified in our study can be considered as appropriate candidate genes affecting neuroticism by within-gene polymorphisms.

For the genes identified in this study, we obtained additional information. For all these genes, we confirmed the independence of the association signal from the effect of the outside SNPs. We found 23 genes associated with neuroticism when using protein-coding SNPs. Three of them, *ANKS1B*, *DEPTOR* and *PANK4,* were identified using only a set of nonsynonymous variants; and nine genes, using both exonic and nonsynonymous variants. Interestingly, for seven of these nine genes, *C12orf49, Me3, MUC22, MYO15A, RPP21, SLC25A37* and *TRIM39-RPP21*, the association signals obtained using nonsynonymous variants were higher than using all exonic variants. The VEP and FATHMM-XF methods detected 18 SNPs in 16 genes as potentially pathogenic variants in accordance with at least one algorithm (Supplementary Table 12). For three of these SNPs (rs10507274, rs35755513, and rs6986), an association with neuroticism had been detected previously ^6,13,20–22^. Several variants have been described as being associated with different behavioral or psychiatric traits such as smoking initiation ^23^, worry ^13^, irritable mood ^20^, feeling guilty ^20^, depression ^13^ (Supplementary Table 16). These results can explain the association of some of the identified genes with neuroticism by the contribution of the potentially pathogenic variants.

For 21 of 23 genes identified using protein-coding SNPs, we found indications of their association with behavioral, cognitive, social or psychiatric traits in the GWAS catalog (Supplementary Table 17). Seven genes, *CYP21A1, MUC22, NOTCH4, RPP21, TAP2, TRIM31* and *TRIM39-RPP21*, were located at the depression loci described by Wray et al. ^24^ (Supplementary Table 18).

Among 167 genes associated with neuroticism using non-coding SNPs, we have found six genes (*PTCH1, FAM120A, ZDHHC5, SNCA, ASCC3* and *MAD1L1*), whose within-gene variation may control the expression of the same genes. For the other eight genes (*MAPT*, *YLPM1*, *STAG1*, *ARHGAP27*, *SETD1A*, *CLUH*, *ZCCHC14* and *GGT7*), we showed the pleiotropy of their within-gene variation with the transcription of adjacent genes (Supplementary Table 13).

In conclusion, we have identified 190 genes associated with neuroticism, 38 of them are new. We have demonstrated that the genes confirmed in our study are more appropriate candidates for neuroticism than the non-confirmed known genes. The majority of the new genes are known as associated with different behavioral, cognitive, social or psychiatric traits. A substantial number of the new genes is represented by the genes identified using the protein-coding SNPs. Many of these genes include potentially pathogenic variants.

## Materials and methods

### Summary statistics

For analysis of neuroticism, we used freely available summary statistics obtained for 10 847 151 genetic variants from a sample of 380 506 participants of the UK Biobank project (https://ctg.cncr.nl/software/summary_statistics). Neuroticism was measured using the Eysenck Personality Questionnaire, Revised Short Form ^25^, consisting of 12 dichotomous items (for details, see Nagel et al. ^20^).

### Gene-based association analysis

The gene-based association analysis was performed using three methods: SKAT-O, PCA and ACAT implemented in the sumFREGAT package ^26^. Then the results of these methods were combined using ACAT ^27^. Correlations between the genotypes of variants within each gene were estimated using the genotypes of about 265 thousand UK Biobank participants.

The results of genome wide gene-based association analysis are freely available in ZENODO database (https://zenodo.org/record/3888340#.XuDTQGlS_IU).

### Regions of interest

SNPs were annotated to genes based on dbSNP version 135 SNP locations and NCBI 37.3 gene definitions. We restricted our gene-based analysis to protein-coding genes. SNPs were annotated to a gene if they were located between its transcription start and stop sites. We used 1000 Genomes functional annotations for genetic variants to determine nonsynonymous, coding and non-coding variants.

We considered four scenarios differing from one another by a set of SNPs whose combined effect was tested by the gene-based association analysis:

1. all SNPs located between the transcription start and stop positions of a gene;
2. SNPs defined as non-coding (introns, 5’UTR and 3’UTR) ones;
3. SNPs defined as exonic (coding) ones;
4. nonsynonymous substitutions.

Two last sets belong to the protein-coding sets.

### Polygene pruning

The purpose of this procedure was to reduce the influence of association signals located outside the region of interest by excluding SNPs that are in high LD with more significant SNPs outside the region. Here we call this exclusion procedure ‘polygene pruning’ to distinguish it from the classical LD pruning and to emphasize the polygenic nature of association signals of excluded SNPs. See Supplementary Methods for a detailed description of the procedure.

We performed polygene pruning only for genes with p < 2.5×10^−5^ demonstrated with at least one of the gene-based methods.

After polygene pruning and re-analysis, genes were considered as identified at p-value < 2.5×10^−5^ in a combined ACAT test.

### Coding variant effect prediction

We considered only protein-coding SNPs with the p-value < 0.05. We used the FATHMM-XF method ^28^ and three prediction methods (SIFT, PolyPhen and CADD) implemented in the Ensembl Variant Effect Predictor (VEP) tool ^29^ to predict the impact of coding SNPs. We defined the status of a variant as pathogenic if at least one of the methods indicated this status: the SIFT p-value < 0.05 (the corresponding qualitative evaluation is “deleterious/deleterious low confidence”), the PolyPhen p-value > 0.5 (“possibly/probably damaging”), the FATHMM-XF p-value > 0.5 (“pathogenic”) and the CADD phred-like score > 20.

### Tissue specificity and gene set enrichment analysis

Gene set enrichment and tissue analyses were carried out with GENE2FUNC implemented in FUMA ^30^ (http://fuma.ctglab.nl/). We used genes identified in the gene-based analysis after polygene pruning as input data.

Enrichment of the identified genes in biological pathways and functional categories was tested against the gene sets obtained from the Gene Ontology (GO) Biological Process and GO Cellular Component. In addition, we tested sets of GWAS catalog-reported genes. Hypergeometric tests were performed to check if the genes of interest were overrepresented in any of the pre-defined gene sets. We used FDR (Benjamini–Hochberg method) for multiple testing correction.

The GTEx v8 54 tissue type data set was used for tissue specificity analysis. A set of the input genes was tested against each of the sets of differentially expressed genes using a hypergeometric test and Bonferroni multiple testing correction.

Statistical significance was determined at an adjusted p-value ◻<◻0.05.

### SMR/HEIDI analysis

Summary data-based Mendelian Randomization (SMR) analysis followed by the Heterogeneity in Dependent Instruments (HEIDI) test ^31^ was used to study potential pleiotropic effects of genes identified using non-coding variants on gene expression levels in different tissues.

SMR analysis provides evidence for pleiotropy (the same locus is associated with two or more traits). It cannot define whether traits in a pair are affected by the same underlying causal polymorphism, but this is done by a HEIDI test, which distinguishes pleiotropy from linkage disequilibrium. It should be noted that SMR/HEIDI analysis does not identify which allele is causal, nor can it distinguish pleiotropy from causation.

Summary statistics for gene expression levels were obtained from Westra Blood eQTL (peripheral blood, http://cnsgenomics.com/software/smr/#eQTLsummarydata) ^32^ and the GTEx version 7 database (53 tissues, https://gtexportal.org) ^33^. We used only blood and tissues related to the brain (14 tissues from GTEx v7). The full list of tissues used in SMR/HEIDI analysis can be found in Supplementary Table 19.

We used the SMR/HEIDI test ^31^ as implemented in the GWAS-MAP platform ^34^. See Supplementary Methods for details.

**Tools used in this study:**

UK biobank project https://ctg.cncr.nl/software/summary_statistics
SumFREGAT:: https://cran.r-project.org/web/packages/sumFREGAT/index.html

1000 Genomes functional annotations for genetic variants:

http://ftp.1000genomes.ebi.ac.uk/vol1/ftp/release/20130502/supporting/functional_annotation/filtered/
FUMA: http://fuma.ctglab.nl/
GWAS catalog: https://www.ebi.ac.uk/gwas/
FATHMM-XF: http://fathmm.biocompute.org.uk/fathmm-xf/about.html
VEP: https://www.ensembl.org/info/docs/tools/vep/index.html
Westra Blood eQTL: http://cnsgenomics.com/software/smr/#eQTLsummarydata
GTEx version 7 database: https://gtexportal.org

## Acknowledgements

**We thank Dr. Yurii Aulchenko and Dr. Gulnara Svishcheva for useful discussion, and Anatoly Kirichenko for technical support.** This work was supported by the Russian Foundation for Basic Research (20-04-00464), and Federal Agency of Scientific Organizations via the Institute of Cytology and Genetics (project 0324-2019-0040-C-01/АААА-А17-117092070032-4). Development of GWAS-MAP platform was supported by the Russian Ministry of Education and Science under the 5-100 Excellence Program.

## Author contributions

All authors contributed to the study conception and design. Material preparation and analysis were performed by Nadezhda Belonogova, Irina Zorkoltseva and Yakov Tsepilov. The first draft of the manuscript was written by Tatiana Axenovich and all authors commented on previous versions of the manuscript. All authors read and approved the final manuscript.

## Conflict of interest

The authors declare that they have no conflict of interest.

## Notes

### Competing Interest Statement

The authors have declared no competing interest.

