## Supplementary material for "Gene-based association analysis identified 190 genes with polymorphisms affecting neuroticism": Supplimental Methods and Tables: Supplementary Methods.pdf

### *Polygene pruning*

The purpose of this procedure was to weaken the influence of association signals located outside the analyzed region by excluding SNPs that are in high LD with more significant SNPs outside the region.

The procedure is as follows:

1. For each SNP with p-value < 0.0001 (index SNP), we formed a list of linked SNPs (clump) from all other SNPs that are in high LD (estimated with 1000G, $r^2 > 0.5$), located within 5 000 kb from the index SNP and having 0.05 > p-value > p-value of the index SNP.
   For clumping, we used PLINK 1.9 with the following options:
   --clump-p1 0.0001
   --clump-p2 0.05
   --clump-r2 0.5
   --clump-kb 5 000
   --clump-range-border 5 000
   --clump-allow-overlap
   --clump-best
   With the --clump-best option, PLINK does not use greedy algorithms and considers all SNPs when constructing each clump. This allowed us to obtain the most complete clumps for all index SNPs. However, some of such clumps contained SNPs with a lower p-value than that of its index SNP. An additional step was to filter these SNPs out of the clumps.
2. For each gene, we formed a list of index SNPs located within or close to the gene (within 5 000 kb from its border). Then we subdivided the list of index SNPs into two sets: one for those located inside and one for those located outside the analyzed region (defined according to one of three types of analysis described in 'Regions of interest' section).
3. For each analyzed region, we removed SNPs if they are in the clumps of index SNPs outside the analyzed region.
4. We re-analyzed the region using the SNPs that remained.

### *SMR/HEIDI analysis*

SMR/HEIDI analysis was conducted as described by Zhu et al. (2016).

HEIDI statistics was calculated as $T_{HEIDI} = \sum_i^m z_{d(i)}^2$, where $m$ is the number of SNPs selected for analysis, $z_{d(i)} = {d_i}/{SE_{(d_i)}}$ and $d_i = \beta_{SMR_i} - \beta_{SMR\ (lead\ SNP)}$.

SNP selection was performed as follows:

1) We defined a set of eligible markers within ±250 kb from the lead SNP in the primary GWAS, which had $\chi^2 > 10$ in the primary GWAS, and for which the results were reported in the secondary GWAS;
2) Created an empty "target" and an empy "rejected" SNP set;
3) From the primary GWAS, selected the SNP with the lowest *p*-value;
4) If this SNP had $r^2 > 0.9$ with any SNP in the target SNP set, we added it to the "rejected" set. The LD matrix ($r^2$) was computed with PLINK 1.9 (https://www.cog-genomics.org/plink2) using 1 000 Genomes data for 503 European individuals (http://www.internationalgenome.org/data/);
5) Otherwise, it was added to the "target" set;
6) The procedure was repeated from step 3 until either eligible SNP set was exhausted or the "target" set had 20 SNPs. If we could not select at least three SNPs, no test was performed.

Analysis was conducted using Python 3.5 as the main programming language.

We selected the genes that were significant in gene-based analysis after pruning using non-coding intronic variants only (164 genes in total). Further, we filtered genes if the p-value for the most associated SNP within the gene (top SNP) was more than $5\times10^{-8}$. It should be noted that the SMR/HEIDI procedure searches for overlapping between both GWAS in analysis. The top SNP for neuroticism was not always present in the expression GWAS panel; in that case, we searched for the second SNP most associated with neuroticism that was also present in the expression GWAS panel (Proxy top SNP). If there was no Proxy top SNP with $r^2 \geq 0.8$ with the top SNP, we removed this locus from analysis. We considered a locus pleiotropic with expression level if the p-values of association for the Proxy top SNP in both neuroticism and expression level GWAS were significant genome-wide (p-value $< 5\times10^{-8}$). The significance threshold for HEIDI tests was set at p-value = 0.001 ($p < 0.001$ corresponds to the rejection of the pleiotropy hypothesis).

Zhu Z, Zhang F, Hu H, Bakshi A, Robinson MR, Powell JE, et al. Integration of summary data from GWAS and eQTL studies predicts complex trait gene targets. Nat Genet. 2016;48(5):481-7.
